# Consolidation Separates Implicit and Explicit Components of Compound Motor Memories

**DOI:** 10.64898/2026.04.15.718660

**Authors:** Adith Deva Kumar, Adarsh Kumar, Neeraj Kumar

## Abstract

Motor adaptation involves the parallel operation of implicit recalibration and explicit re-aiming processes. During naturalistic learning, these systems interact, producing compound behavioral outputs that reflect their combined contributions. It remains unclear whether simultaneously engaged implicit and explicit processes form a single unified representation, or generate parallel memory representations that are merely co-expressed, and how consolidation transforms such representations. We addressed these questions across three visuomotor adaptation experiments (*n* = 120), in which the implicit process was engaged via gradual cursor rotation and the explicit process via target jump, by systematically manipulating the sequence of learning and the timing of expression. Immediately after learning, behavior reflected an inflexible, integrated memory that could not be decomposed by changing task demands. Following 24-hour consolidation, however, expression became component-selective, with implicit or explicit contributions retrieved in response to task demand. This reorganization had direct consequences on relearning, producing facilitation when the expressed and relearned components matched and interference when they mismatched. Moreover, when implicit adaptation was stabilized prior to compound learning, consolidation preserved the updated state rather than the original implicit representation. Together, these findings demonstrate that consolidation does not merely stabilize compound motor memories. Instead, it actively reorganizes them, transforming the initially integrated representations into independent, context-dependent components.

## Introduction

Our motor system learns, adapts, and stores multiple memories of the movements that guide our future actions in our day-to-day life (Krakauer & Mazzoni, 2011; Wolpert et al., 2011). Motor behavior is shaped by distinct learning processes that operate across multiple timescales and levels of control (Diedrichsen et al., 2010; Huang et al., 2011). A central question in the domain of motor neuroscience is how these processes contribute during learning and, importantly, how their contributions are stored and expressed over time. Work over the past two decades has established that motor adaptation is supported by the interaction of distinct learning processes, rather than a single learning mechanism (Haith & Krakauer, 2013; Lee & Schweighofer, 2009; McDougle et al., 2015; Smith et al., 2006; Taylor & Ivry, 2011). These processes are sensitive to different sources of error, where the slow, implicit recalibration process is usually driven by the sensory prediction error (Kawato, 1999; Mazzoni & Krakauer, 2006; Taylor et al., 2010; Tseng et al., 2007; Wolpert & Miall, 1996), while the fast, strategic, explicit process is driven by task error (Haith & Krakauer, 2013; Körding & Wolpert, 2004; Leow et al., 2020; Morehead et al., 2015; Taylor et al., 2014). In naturalistic learning, these error signals co-occur, engaging both implicit and explicit systems simultaneously (Krakauer et al., 2019; Taylor et al., 2014). Crucially, these systems do not operate independently but interact during adaptation (Albert et al., 2022; Bond & Taylor, 2015; Haith & Krakauer, 2013). When both are engaged, motor output reflects their combined influence rather than the isolated contribution of either system (McDougle et al., 2015; Taylor et al., 2014). For instance, during visuomotor rotation tasks, participants simultaneously develop implicit recalibration and explicit re-aiming strategies that jointly determine reaching direction (Bond & Taylor, 2015; Mazzoni & Krakauer, 2006). When subsequently asked to perform the learned movement without constraints, behavior reflects this compound expression of both learning mechanisms (Morehead et al., 2017).

Importantly, dissociations between implicit and explicit components typically require specialized probe conditions designed to isolate one process (Kim et al., 2019; Leow et al., 2017; Morehead et al., 2017; Vaswani et al., 2015). Under unconstrained conditions, behavior reflects an integrated memory state in which both contributions are merged (Haith & Krakauer, 2013; Taylor et al., 2014). This suggests that, when no constraints were imposed during expression of the adapted behavior, the motor system expresses an integrated memory of both learning mechanisms while present in a working memory buffer (Christou et al., 2016). This observation raises a fundamental question about memory architecture: when multiple learning systems interact, do they create a single, unified representation, or do they form parallel traces that are merely expressed concurrently? Critically, this question can be asked at two time points: immediately after learning and after consolidation has had time to act.

This question becomes particularly consequential when considering memory consolidation, the process by which labile memories are stabilized and transformed over time (McGaugh, 2000; Müller & Pilzecker, 1900; Squire & Alvarez, 1995). Consolidation is known to strengthen motor memories against interference (Dudai, 2004; Krakauer & Shadmehr, 2006; Robertson et al., 2004; Shadmehr & Brashers-Krug, 1997), but how it operates on memories formed through multi-system interaction remains unexplored. Critically, implicit and explicit learning engage distinct neural circuits. The implicit adaptation relies primarily on the cerebellar error-correction circuits (Butcher et al., 2017; Flament et al., 1996; Hadjiosif et al., 2014; Izawa et al., 2012; Smith & Shadmehr, 2005; Tseng et al., 2007), whereas the explicit strategy engages prefrontal and parietal networks (Anguera et al., 2010; Benson et al., 2011; Clower et al., 1996; Della-Maggiore et al., 2004; Doyon et al., 2009; A. Kumar et al., 2020; Seidler et al., 2012). This neural dissociation raises two competing possibilities about how consolidation might transform these memories. One possibility is that consolidation simply stabilizes the integrated state, preserving the compound memory as a unified representation. If this were the case, post-consolidation expression should remain integrated across task contexts, and relearning should be similar across conditions. An alternative possibility is that consolidation actively reorganizes memory architecture, separating initially integrated representations into their constituent components. This view predicts that after consolidation, implicit and explicit components should become independently accessible, with expression selectively gated by task demands. Furthermore, if components become separable, their expression should determine subsequent learning: facilitation when the expressed component matches new task demands and interference when it mismatches.

The present study tests these competing predictions by asking three specific questions: First, do compound memories formed through simultaneous implicit-explicit engagement consolidate as integrated wholes or separable components? Second, do individual components maintain independent stability across consolidation when isolated from competition? Third, does the sequence of learning alter component strength and the nature of consolidation?

To answer these questions, we conducted three visuomotor adaptation experiments that manipulated the timing, sequence, and context of implicit and explicit learning. In all experiments, implicit learning was engaged through gradual cursor rotation, which preferentially drives recalibration based on sensory prediction error without conscious awareness (Kagerer et al., 1997; Klassen et al., 2005; Tseng et al., 2007). Explicit learning was engaged through target jumps (30° displacement at movement onset), which require conscious re-aiming and elevate reaction times – a hallmark of strategic control (Fernandez-Ruiz et al., 2011; Haith et al., 2015; Taylor et al., 2014). This combination allowed us to independently manipulate and measure each learning system’s contribution. Overall findings suggest that immediate expression reflects a unified compound memory, whereas consolidation reorganizes it into dissociable components whose later expression depends on task demands.

## Methods

### Subjects

The study involved 120 healthy right-handed participants with a mean age of 22.67 ± 3.38 years (61 females). All participants reported no neurological disorders, psychological conditions, cognitive impairments, or orthopedic injuries, and they all had normal or corrected-to-normal vision. Prior to the experiment, participants were naive to the study’s objective, provided written informed consent to participate, and their handedness was assessed using the Edinburgh Handedness Inventory (Oldfield, 1971). Upon successful completion of the task, participants received a monetary reward for their time and participation. This study was reviewed and approved by the Institutional Ethical Committee.

### Experimental setup and trial structure

Participants sat in a completely dark, sound-attenuated room on an adjustable-height chair, facing a virtual reality setup with a digitizing tablet (GTCO CalComp). They performed a planar point-to-point reaching task using a handheld stylus on the tablet. A semi-silvered mirror was mounted horizontally at eye level beneath a high-definition display (1920 × 1080 pixels, 120 Hz), serving as the viewing platform, and preventing participants from directly seeing their arm movements over the tablet positioned below (fig.1A). Hand movement data were sampled at 120 Hz.

**Figure 1:**
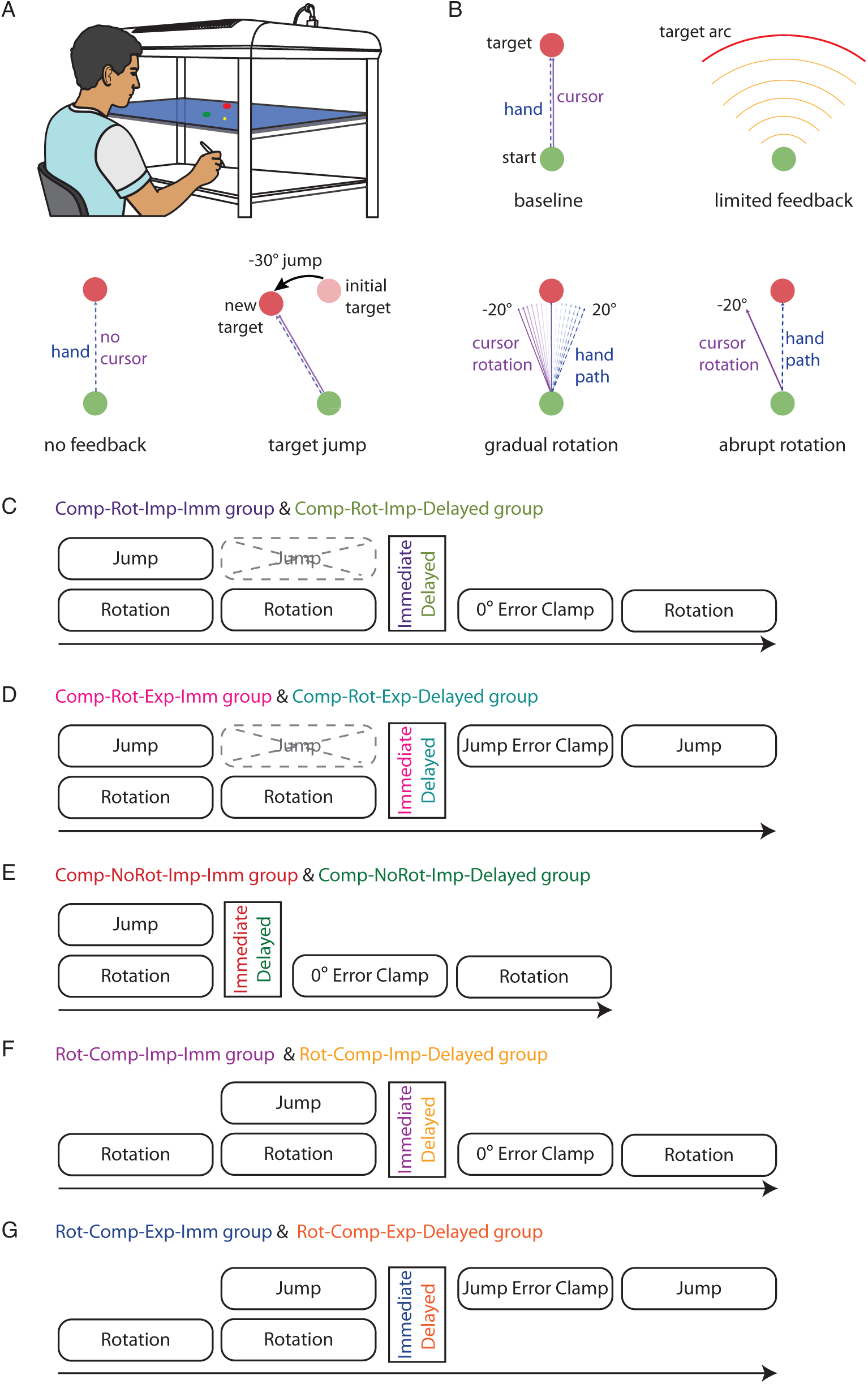
A. A virtual-reality reaching setup was used, in which an HD monitor projected the screen onto a semi-silvered mirror positioned at eye level, blocking direct vision of the hand. Participants made planar reaches on a digitizing tablet placed beneath the mirror using a handheld stylus. The mirror display presented the start position, targets, and a cursor that provided veridical feedback of the stylus location. B. Schematic illustration of the task manipulations used across experiments. Baseline: Participants reached from a start position to a stationary target with veridical cursor feedback. Limited-feedback trials: Only endpoint cursor information was provided along a target arc. No-feedback trials: The cursor was removed, requiring it to reach directly to the visible no-jump target. Target-jump trials: The target was displaced to –30° at movement onset, requiring rapid re-aiming to the new location. Gradual rotation: Cursor feedback was rotated incrementally from 0° to −20° over trials. Abrupt rotation: A −20° cursor rotation was introduced suddenly. C. Comp-Rot-Imp-Imm group: Perturbation schedule showing baseline, 200-trial target-jump+gradual-rotation learning block, 100-trial rotation-only block, five zero-clamp retention trials, and a 100-trial abrupt-rotation relearning block (violet). Comp-Rot-Imp-Delayed group: Identical task structure to Comp-Rot-Imp-Imm group, but retention (zero clamp) and relearning (abrupt rotation) were completed after a 24-hour delay (olive). D. Comp-Rot-Exp-Imm group: Same learning structure as Comp-Rot-Imp-Imm group, followed by five jump-clamp retention trials and a 100-trial jump-only relearning block (pink). Comp-Rot-Exp-Delayed group: Same structure as Comp-Rot-Exp-Imm group, with retention (jump clamp) and jump-only relearning administered after a 24-hour delay (cyan). E. Comp-NoRot-Imp-Imm group: Participants completed baseline and a single 200-trial target-jump+gradual-rotation block, followed immediately by zero-clamp retention trials and a 100-trial abrupt-rotation relearning block (no rotation-only block provided, red). Comp-NoRot-Imp-Delayed group: Same perturbation schedule as Comp-NoRot-Imp-Imm group, but retention (zero clamp) and relearning (abrupt rotation) occurred after a 24-hour delay (green). F. Rot-Comp-Imp-Imm group: Participants completed a 200-trial gradual-rotation block followed by a 200-trial target-jump+gradual-rotation block, then immediate zero-clamp retention trials and a 100-trial abrupt-rotation relearning block (purple). Rot-Comp-Imp-Delayed group: Same learning structure as Rot-Comp-Imp-Imm group, with retention (zero-clamp) and abrupt-rotation relearning performed after a 24-hour delay (yellow). G. Rot-Comp-Exp-Imm group: Same learning sequence as Rot-Comp-Imp-Imm group, followed by immediate retention using jump-clamp trials and a 100-trial jump-only relearning block (blue). Rot-Comp-Exp-Delayed group: Same learning structure as Rot-Comp-Exp-Imm group, with retention (jump-clamp) and jump-only relearning delivered after a 24-hour delay (orange).

**Table 1:** Experimental design and group structure. The table summarizes the sequence of learning sessions, timing of expression, retention tests, and relearning conditions for each group. Group names encode the order and type of learning phases, the component assessed during retention, and the timing of expression. Retention was evaluated using error-clamp trials designed to isolate either implicit or explicit contributions, and relearning was assessed under rotation-only or jump-only conditions. Note: CCW = counterclockwise.

| Experiment | Group | Learning Session-1 | Learning Session-2 | Timing of Expression | Retention Test | Relearning Session |
| --- | --- | --- | --- | --- | --- | --- |
| 1 | Comp-Rot-Imp-Imm | Compound learning **(Comp)**<br>[Target jumps 30° CCW with Gradual rotation of 20° CCW] | Rotation-only **(Rot)**<br>[Target jump removed] | Immediate **(Imm)** | 0° error-clamp **(Imp)**<br>[Cursor clamped to Target] | Rotation-only<br>[20° CCW] |
|  | Comp-Rot-Imp-Delayed |  |  | 24h later **(Delayed)** |  |  |
|  | Comp-Rot-Exp-Imm |  |  | Immediate **(Imm)** | Jump error-clamp **(Exp)**<br>[Cursor clamped to jumping Target] | Jump-only<br>[30° CCW] |
|  | Comp-Rot-Exp-Delayed |  |  | 24h later **(Delayed)** |  |  |
| 2 | Comp-NoRot-Imp-Imm | Compound learning **(Comp)**<br>[Target jumps 30° CCW with Gradual rotation of 20° CCW] | No session **(NoRot)** | Immediate **(Imm)** | 0° error-clamp **(Imp)**<br>[Cursor clamped to Target] | Rotation-only<br>[20° CCW] |
|  | Comp-NoRot-Imp-Delayed |  |  | 24h later **(Delayed)** |  |  |
| 3 | Rot-Comp-Imp-Imm | Gradual rotation **(Rot)**<br>[20° CCW] | Compound learning **(Comp)**<br>[Target jumps 30° CCW with rotation of 20° CCW] | Immediate **(Imm)** | 0° error-clamp **(Imp)**<br>[Cursor clamped to Target] | Rotation-only<br>[20° CCW] |
|  | Rot-Comp-Imp-Delayed |  |  | 24h later **(Delayed)** |  |  |
|  | Rot-Comp-Exp-Imm |  |  | Immediate **(Imm)** | Jump error-clamp **(Exp)**<br>[Cursor clamped to jumping Target] | Jump-only<br>[30° CCW] |
|  | Rot-Comp-Exp-Delayed |  |  | 24h later **(Delayed)** |  |  |

The cursor was denoted by a yellow dot with a diameter of 0.3 cm, while the start point was a blue circle with a diameter of 0.6 cm presented at the bottom center of the screen. At the start of each trial, participants were required to bring the cursor to the start point and stay there for a duration of 0.5 seconds, after which the start point turned green (go cue), and participants were required to move to the red target circle (0.6 cm in diameter) presented to them. A single target was presented at an angle of 90 degrees, 12 cm straight from the start point. Participants were instructed to reach the target within 1.5 seconds and maintain their position until an auditory cue signaled the end of the trial. After every trial, they received visual feedback in the form of a numerical score reflecting their accuracy (0-10 points, depending on proximity to the target), along with a rating of movement velocity (too fast, good, or too slow). Points awarded during the task were not used for data analysis and did not affect participants’ compensation. Cursor feedback was presented throughout the entire movement and was either veridical or distorted relative to the hand’s motion, depending on the trial type.

After instructions on the task and a brief familiarization with the experimental setup, and a few practice reaching trials, participants performed the visuomotor task under different trial conditions, such as baseline, rotation, target jump, catch, and error-clamp. During baseline trials, participants were asked to make ballistic movements towards the presented target with veridical cursor feedback throughout the movement. In target jump trials, the target shifted to a new location as soon as the cursor moved 0.5 cm away from the center of the start circle. This shift in the target was executed by removing the initially presented target (original target) and instantly providing the new target in the jump location, which was always a –30° counterclockwise to the original target (fig.1B). Here, participants were asked to ignore the location of the original target and were asked to make movements towards the new target. During rotation trials, the cursor feedback was gradually rotated to an angle of –20° counterclockwise (ramp of –0.4° per trial followed by a hold phase) to the position of the hand during movement towards the target. Limited feedback catch trials were presented within the rotation block to assess the implicit learning, where the target circle was replaced by a red target arc, and the cursor appeared as a yellow semi-circular arc that expanded in diameter as participants moved forward. Participants were instructed to stop their movement when the arc reached the target arc, thereby constraining movement magnitude without providing direction-related information during movement. A total of four sub-blocks comprising two limited feedback catch trials were interspersed systematically within the hold phase of the rotation block (fig.1B). During the target jump block, no-jump catch trials were presented where the target would not jump, and the cursor would not be visible while moving to the target. Here, similar to limited feedback catch trials, 8 no-feedback trials were provided, and participants were instructed right before the trials that the target would not jump, and they had to move their hand to the presented target in the absence of any feedback (fig.1B). The error-clamp trials are used to assess the retention. In zero-clamp trials, cursor movement was constrained to 0° error relative to the original target. In jump-clamp trials, the target jumped to –30° at movement onset, and cursor movement was clamped to the jump location (0° error relative to the jumped target). Participants were not informed of the clamp.

### Experiment-1

Experiment 1 was designed to test whether consolidation preserves an integrated memory of simultaneously learned implicit and explicit components, or instead enables their separation into independently retrievable components. Forty-eight participants were assigned to four groups in a 2 × 2 factorial design that varied the timing of retention testing (immediate vs. 24-h delayed) and the component probed during retention (implicit vs. explicit). The groups were: Comp-Rot-Imp-Imm (compound learning, rotation-only session present, implicit probe, immediate), Comp-Rot-Exp-Imm (compound learning, rotation-only session present, explicit probe, immediate), Comp-Rot-Imp-Delayed (compound learning, rotation-only session present, implicit probe, 24-h delay), and Comp-Rot-Exp-Delayed (compound learning, rotation-only session present, explicit probe, 24-h delay).

All groups followed the same initial protocol: after 30 baseline trials, participants performed a compound learning block (200 trials) in which a –30° (counterclockwise) target jump was present on every trial and a –20° (counterclockwise) visuomotor rotation was introduced gradually (–0.4° per trial for the first 50 trials, then held constant for the remaining 150 trials). Participants were explicitly told that the target would jump on each trial and were instructed to aim the movement to the jumped target. During the hold phase of this block, four no-jump catch sub-blocks (2 trials each) were presented, where participants were asked to reach directly to the original target (no jump, no cursor feedback). These catch trials measured the evolving implicit aftereffect while the explicit strategy was simultaneously engaged.

Immediately after compound learning, participants underwent a rotation-only session (100 trials) in which the target jump was removed, but the rotation remained. Four limited-feedback catch sub-blocks (2 trials each) were interspersed to assess adaptation under pure implicit demand. This rotation-only session was designed to probe the immediate flexibility of the compound memory. If implicit and explicit components were already stored as independent representations before consolidation, removing the explicit demand should have produced an immediate shift to pure implicit adaptation. Conversely, if the memory was integrated, behavior would reflect carryover of the compound policy, revealing that the pre-consolidation memory state could not be flexibly decomposed.

After the rotation-only session, retention was tested using clamp trials that isolated either the implicit or the explicit component. For groups testing the implicit component (Comp-Rot-Imp-Imm and Comp-Rot-Imp-Delayed), participants completed five zero-clamp trials (cursor clamped to the original target). They were instructed that the target would not jump and were asked to reproduce the hand movement they had executed at the end of the rotation-only block (the last block before retention). For groups testing the explicit component (Comp-Rot-Exp-Imm and Comp-Rot-Exp-Delayed), participants completed five jump-clamp trials (target jumped to –30°, cursor clamped to the jump location). They were instructed to reproduce the hand movement they had used at the end of the compound learning block (the block in which the target had been jumping). In all cases, participants were unaware that the cursor was clamped.

Immediately after retention (for the immediate groups) or 24 h later (for the delayed groups), participants performed a relearning block of 100 trials. The relearning task matched the component that had been probed during retention: groups that had been tested on the implicit component (Comp-Rot-Imp-Imm and Comp-Rot-Imp-Delayed) relearned an abrupt –20° rotation (no target jump); groups that had been tested on the explicit component (Comp-Rot-Exp-Imm and Comp-Rot-Exp-Delayed) relearned the –30° target jump alone (no rotation). Limited-feedback catch trials were again interspersed during the relearning session.

This design allowed us to contrast the pre-consolidation state (observed during the rotation-only session) with the post-consolidation state (observed in the clamp retention tests). The critical prediction was that if consolidation actively separates the components, the delayed groups would show cue-dependent expression – implicit behavior when cued for rotation, explicit behavior when cued for jump—even though immediately after learning the system had been unable to flexibly decompose the compound memory (as evidenced by carryover during the rotation-only session). Relearning performance then provided an additional test of whether the expressed component determined subsequent facilitation or interference.

### Experiment-2

Experiment 2 tested whether the implicit component acquired during compound learning can be accessed directly and remains stable across consolidation, without the intervening rotation-only session used in Experiment 1. In Experiment 1, the rotation-only session revealed that the pre-consolidation memory state was inflexible, but it also provided participants with a block of pure implicit-only experience before retention. This raised the possibility that the implicit component expressed during retention might have been shaped or strengthened by that additional exposure. Experiment 2, therefore, omitted the rotation-only session, allowing us to isolate the implicit component immediately after compound learning and to assess its persistence over 24 hours. Twenty-four participants were assigned to two groups that differed only in the timing of retention testing: Comp-NoRot-Imp-Imm (compound learning, no rotation-only session, implicit probe, immediate), and Comp-NoRot-Imp-Delayed (compound learning, no rotation-only session, implicit probe, 24-h delay). Both groups completed the same initial learning phase. After a 30-trial baseline, participants performed a compound learning block of 200 trials identical to that used in Experiment 1: a –30° target jump on every trial and a gradual –20° rotation (–0.4° per trial for 50 trials, then held constant for 150 trials). Participants were instructed to reach quickly and straight to the jumped target. During the hold phase, catch trials measured the ongoing implicit aftereffect during compound learning, while the explicit strategy was still active.

Critically, after compound learning, participants did not receive a rotation-only session. Instead, they proceeded directly to a retention test consisting of five zero-clamp trials (cursor clamped to move straight to the original target). The instruction was: “The target will not jump. Aim straight to the displayed target.” This instruction was identical to that used during the catch trials, but now it was given after the compound learning block had ended, and no explicit strategy was being maintained. Thus, the zero-clamp trials measured the residual implicit aftereffect in the absence of any ongoing explicit engagement or any intervening implicit-only experience.

Retention was tested either immediately after compound learning (Comp-NoRot-Imp-Imm) or after a 24-hour delay (Comp-NoRot-Imp-Delayed). Following retention, both groups completed a relearning block of 100 trials in which an abrupt –20° rotation was introduced (no target jump), with interspersed limited-feedback catch trials.

If the implicit component is inherently present and stable, it should be measurable in the zero-clamp retention test regardless of whether the rotation-only session was experienced. Moreover, its magnitude should not differ between immediate and delayed testing, indicating that consolidation does not degrade the implicit memory. Conversely, if the rotation-only session in Experiment 1 had artificially enhanced or altered the implicit component, then the implicit aftereffect measured in Experiment 2 (without that session) would be weaker, or might decay over 24 hours. By removing the rotation-only block, Experiment 2 thus provided a critical test of the stability of the implicit component across consolidation.

### Experiment-3

This experiment tested how learning history affects component strength and updatability. In Experiments 1 and 2, implicit and explicit components were acquired simultaneously, leaving open the question of whether prior stabilization of one component would alter its strength or its capacity to be updated by subsequent experience. Here, we reversed the learning sequence: participants first learned the implicit rotation in isolation, allowing it to stabilize before explicit demands were introduced. This design addressed two specific questions. First, does stabilization of implicit adaptation produce a stronger implicit component that resists decay when an explicit strategy is later superimposed? Second, when the explicit strategy alters the sensorimotor relationship (by adding a target jump to an already-adapted implicit system), does the implicit component update, and if so, which version-the original implicit-only state or the updated recalibrated state—does consolidation preserve?

Forty-seven participants were randomly assigned to four groups in a 2 × 2 factorial design that varied the timing of retention testing (immediate vs. 24-h delayed) and the component probed during retention (implicit vs. explicit). The groups were: Rot-Comp-Imp-Imm (rotation-only session, compound learning, implicit probe, immediate), Rot-Comp-Exp-Imm (rotation-only session, compound learning, explicit probe, immediate), Rot-Comp-Imp-Delayed (rotation-only session, compound learning, implicit probe, 24-h delay), and Rot-Comp-Exp-Delayed (rotation-only session, compound learning, explicit probe, 24-h delay). All groups followed the same initial two-phase learning protocol. After a 30-trial baseline, participants first completed a rotation-only learning block of 200 trials, during which a –20° visuomotor rotation was gradually introduced (–0.4° per trial for the first 50 trials, then held constant). Participants were not informed of the rotation. During the hold phase, four limited-feedback catch sub-blocks were interspersed to assess the state of implicit adaptation. This block established a stable implicit-only adaptation state.

Immediately after the rotation-only block, participants were instructed that the target would now jump, and they completed a compound learning block of 100 trials in which a –30° target jump was added while the –20° rotation remained active. During this block, four no-jump catch sub-blocks were interspersed. In these catch trials, the target did not jump, and the cursor feedback was withheld; participants were instructed to reach directly to the original target. These catch trials measured the implicit aftereffect during compound learning, after the explicit strategy had been superimposed on the pre-existing implicit adaptation.

Following the two learning blocks, retention was tested using clamp trials that isolated either the implicit or the explicit component, with the instruction tailored to the probe type. For groups testing the implicit component (Rot-Comp-Imp-Imm and Rot-Comp-Imp-Delayed), participants completed five zero-clamp trials (cursor clamped to move straight to the original target). The instruction was: “The target will not jump. Reproduce the hand movement you executed at the end of the rotation-only block, before the target jumps were introduced.” This instruction cued the retrieval of the implicit component as it existed before compound learning. For groups testing the explicit component (Rot-Comp-Exp-Imm and Rot-Comp-Exp-Delayed), participants completed five jump-clamp trials (the target jumped to –30°, and the cursor was clamped to the jump location). The instruction was: “The target will jump. Reproduce the hand movement you executed at the end of the compound learning block.” This instruction cued the retrieval of the compound behavior after the explicit strategy had been superimposed. In all cases, participants were unaware of the cursor clamp.

Retention was tested either immediately after learning (for the immediate groups) or after a 24-hour delay (for the delayed groups). Following retention, participants performed a relearning block of 100 trials matched to the component that had been probed: groups that had been tested on the implicit component (Rot-Comp-Imp-Imm and Rot-Comp-Imp-Delayed) relearned an abrupt – 20° rotation (no target jump); groups that had been tested on the explicit component (Rot-Comp-Exp-Imm and Rot-Comp-Exp-Delayed) relearned the –30° target jump alone (no rotation). Limited-feedback catch trials were interspersed during relearning.

If prior stabilization of implicit learning produces a stronger component, the implicit aftereffect measured during the compound learning catch trials should be larger than that observed in Experiment 1, where implicit and explicit were learned simultaneously. Moreover, if the implicit system remains updatable when explicit strategy alters the sensorimotor relationship, the implicit aftereffect during compound learning should differ from the original rotation-only state – evidence of recalibration driven by sensory prediction errors generated by the superimposed explicit strategy. Finally, comparing the implicit component expressed during zero-clamp retention (cued to the rotation-only block) between immediate and delayed groups tested which version of consolidation preserves: if consolidation stabilizes the most recent functional state, the delayed group should express the updated implicit component (measured during compound learning catch trials) rather than reverting to the original implicit-only state. Relearning performance then provided an additional test: facilitation would occur when the expressed component matched the relearning task, interference when mismatched.

Together, these experiments test a novel theoretical proposition: that consolidation does not merely stabilize motor memories but actively restructures them, transforming initially rigid, integrated representations into flexible, modular components whose expression becomes context-dependent. This reorganization would represent a fundamental principle of memory optimization—the brain’s solution to balancing stability with behavioral flexibility. By revealing how implicit and explicit memories separate and recombine, our findings illuminate basic mechanisms of skill learning, retention, and transfer with implications for rehabilitation, training, and human-machine interaction.

### Data pre-processing and analysis

Hand movement data were analyzed using custom MATLAB scripts. Hand position data were low-pass filtered using a 10 Hz Butterworth filter, and movement speed was derived by differentiating positional data. Movement onset was defined as the first time point at which the cursor crossed the start point circle. Reaction time (RT) was calculated as the interval between target appearance and movement onset. Hand angles were computed as the angle between the line connecting the start circle to the target and the line connecting the start circle to the hand position at peak tangential velocity. For target-jump trials, hand angles were calculated relative to the new target location. All hand angles were baseline corrected. Hand angle and RT data were binned by averaging across one cycle of 5 trials. Catch trials were a pair of two, hence they were binned separately from the learning data. For catch trials, only the last bin of the compound learning block was used for comparison with the retention block to improve robustness and to represent the stable mechanism.

For Experiments 1-3, statistical analyses were performed separately for immediate and delayed retention groups. One-sample t-tests were used to assess aftereffects in catch trials and zero-/jump-clamp trials against zero, quantifying the contribution of implicit adaptation. Paired t-tests compared hand angles between the last learning bin and retention trials to examine immediate expression of learned mechanisms, while unpaired t-tests assessed differences between immediate and delayed groups to determine the effect of consolidation on the expression of implicit and explicit components. Reaction times were similarly analyzed to quantify engagement of explicit strategic mechanisms, and mixed ANOVAs were used to examine session × group interactions when assessing changes across sessions and groups. Where multiple comparisons across sessions or groups were performed, Tukey’s post hoc tests were conducted following ANOVAs to identify pairwise differences. Effect sizes were reported using Cohen’s *d* for t-tests and partial eta-squared (η²p) for ANOVAs.

All analyses were performed using MATLAB and R (v4.3.1). Unless otherwise stated, results are reported as mean ± SD, with 95% confidence intervals provided for all statistical comparisons.

## Data exclusion

Of the 120 recruited participants, data from one participant in the Rot-Comp-Exp-Imm group in Experiment-3 were excluded due to failure to follow task instructions, resulting in a final sample of 119 participants included in the analyses. Across all experiments, 0.38% of trials were excluded due to stylus disconnection or failure to initiate movement. Of the remaining trials, 0.15% were discarded due to extreme hand angles (±50°). All subsequent analyses were conducted on the cleaned dataset following these exclusion criteria.

## Results

Across three experiments, we systematically manipulated the kind of interaction between the implicit (gradual visuomotor rotation-driven recalibration) and explicit (target jump–driven strategic re-aiming) learning mechanisms. This framework allowed us to examine the nature of the memory formed by these processes when engaged simultaneously, and to understand how it changes after consolidation.

In experiment-1, four groups (n=48) were involved, where the following baseline of 30 trials, all participants learned to compensate for both target jump and rotation simultaneously, consisting of 200 trials in session-1. Following that, all 4 groups experienced 100 trials of only rotation in the absence of the target jump in session 2. This was followed by the error-clamp session (5 trials) to test retention of the learning mechanisms engaged previously. In Comp-Rot-Imp-Imm group (zero-error clamp), participants were told that there would be no target jump and were asked to move their hand exactly as they did at the end of Session 2, that is, the rotation alone behavior. In Comp-Rot-Exp-Imm group (jump-error clamp), participants were informed that the target would jump and were asked to move the same way they did at the end of Session 1, that is, the target-jump+gradual-rotation behavior. During the retention session, participants were neither aware nor informed explicitly that the cursor was clamped during these trials. Following the retention trials, participants in Comp-Rot-Imp-Imm group performed a relearning session with the same degree of abrupt rotation, whereas participants in Comp-Rot-Exp-Imm group relearned only the 30° target jump without any rotation. The remaining two groups, Comp-Rot-Imp-Delayed group (zero-error clamp delayed) and Comp-Rot-Exp-Delayed group (jump-error clamp delayed), completed the same retention and relearning sessions after a 24-hour delay (Day 2), mirroring the task structure of Comp-Rot-Imp-Imm group and Comp-Rot-Exp-Imm group, respectively, but testing retention and relearning after consolidation.

### Simultaneous engagement resulted in a compound adaptive mechanism during learning

Following baseline with veridical feedback and no perturbations, a target jump of –30 degrees was introduced along with a gradual cursor rotation of –20 degrees with an increment of –0.4 degrees over 50 trials and a hold of –20 degrees over 150 trials, simultaneously with 200 trials of target jump. Participants were able to compensate for the target jump through explicit strategies during initial exposure trials, as evidenced by a rapid adjustment of hand angle to the introduced jump. Along with it, they also showed a gradual recalibration of their hand movement in response to the introduced gradual rotation, where they eventually reached an angle of around –10 degrees at the end of session 1, compensating for both the jump and the rotation (fig.2A and 2C). Participants from all the groups from experiment 1 were able to adjust hand direction in response to both the perturbations that were introduced together (last bin of session 1, mean ± SD = Comp-Rot-Imp-Imm group: –11.28 ± 4.72; Comp-Rot-Exp-Imm group: –12 ± 2.4; Comp-Rot-Imp-Delayed group: –13.73 ± 3.4; Comp-Rot-Exp-Delayed group: –11.01 ± 3.11). A one-way ANOVA revealed no significant effect of group on adaptation at the end of the compound learning session, F(3, 44) = 1.46, *p* = 0.24, η²p = 0.09, suggesting the learning was the same across these groups.

**Figure 2.**
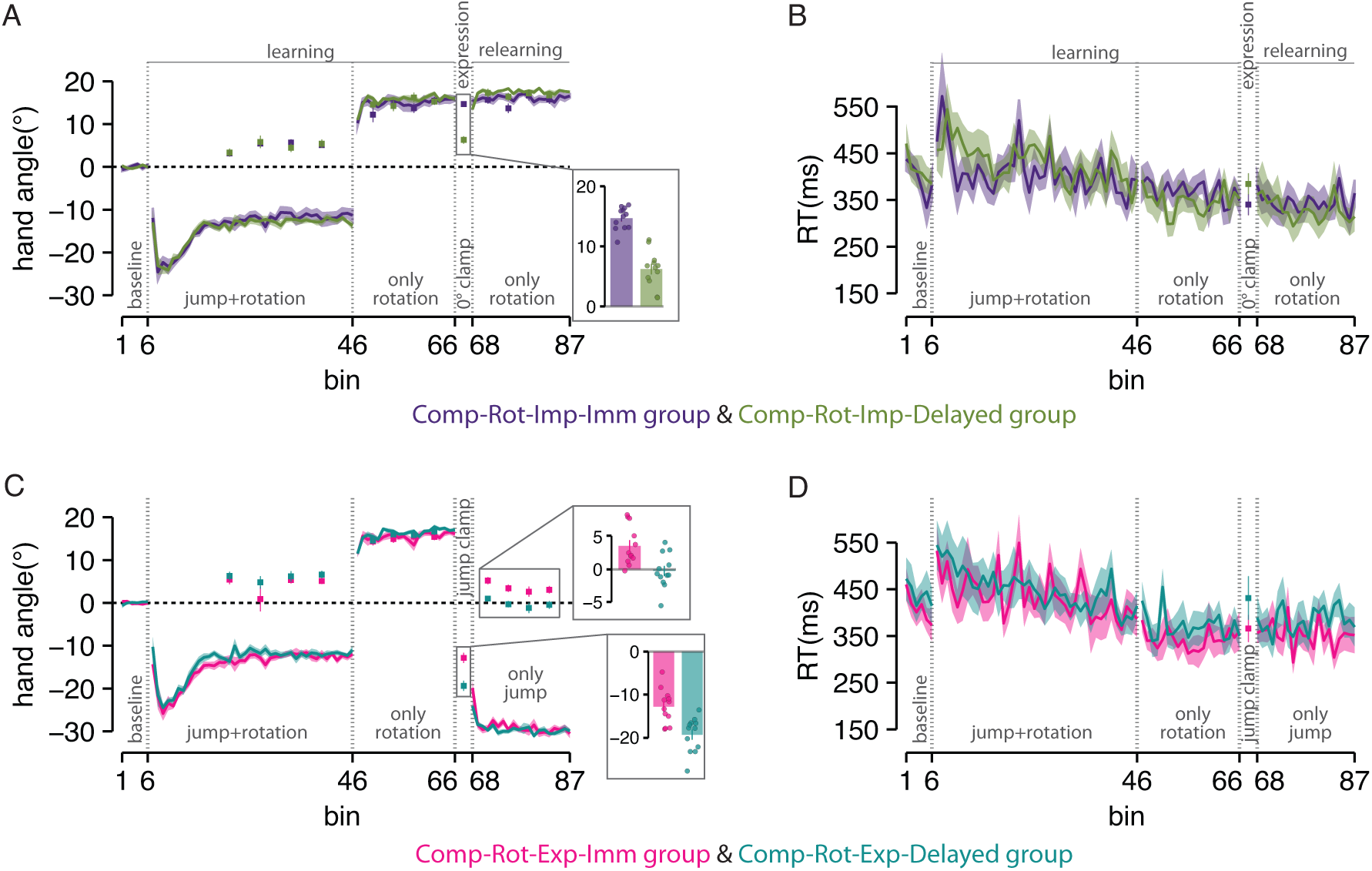
Consolidation alters the expression and relearning of the compound mechanism acquired during combined jump–rotation learning. A, C. Group-averaged, binned hand-angle trajectories relative to the original target. Participants showed smooth acquisition of the compound perturbation (target-jump+gradual-rotation) and exhibited clear aftereffects during no-feedback catch trials, demonstrating engagement of implicit recalibration. In the subsequent rotation-only block, expression of the compound memory persisted immediately, despite removal of the target-jump component. Bar plots in the bottom (inset panel in A and C) compared hand angle between the immediate and delayed groups during the expression session. Immediate groups (Comp-Rot-Imp-Imm: A, violet; Comp-Rot-Exp-Imm: C, pink) expressed the full compound memory during retention (error-clamp) trials. After a 24-hr delay, delayed groups (Comp-Rot-Imp-Delayed: A, olive; Comp-Rot-Exp-Delayed: C, cyan) expressed memory of the individual components rather than the compound behavior: implicit-dominant expression in the rotation-only condition and explicit-dominant expression in the jump-only condition. The top inset panel in C shows the aftereffects observed in the jump-alone relearning session. This shows that implicit aftereffects persist in the immediate group but are absent after consolidation, indicating a shift toward explicit-dominant expression in the delayed group. during the jump-only relearning condition. B, D. Reaction times (group-averaged, binned) increased sharply during compound learning, reflecting explicit strategy use for compensating the target jump, and remained elevated during the transition into the rotation-only block, indicating that the explicit component of the compound mechanism carried over.

We introduced the catch trials during the hold phase of session 1, interspersed to verify the existence and engagement of the implicit mechanism, along with the explicit mechanism for learning target jump. In these catch trials, participants were instructed that the target would not jump and asked to aim at the original target to measure aftereffects of implicit adaptation. Significant aftereffects were observed as hand deviations in the catch trials during the session1 (one sample t-test, Comp-Rot-Imp-Imm group: 4.87 ± 1.99, t(11) = 8.45, *p* < 0.001, 95% confidence interval (CI) = [3.60, 6.14], Cohen’s *d* = 2.43; Comp-Rot-Exp-Imm group: 4.46 ± 3.13, t(11) = 4.94, *p* < 0.001, 95% CI = [2.48, 6.45], Cohen’s *d* = 1.42), suggesting that implicit learning mechanism was active during the learning of session 1 (fig.2A-violet and 2C-pink). Similarly, hand deviation during catch trials when compared with zero yielded significant differences in both the delayed groups during session 1 (one sample t-test, Comp-Rot-Imp-Delayed group: 4.62 ± 2.29, t(11) = 6.97, *p* < 0.001, 95% CI = [3.16, 6.08], Cohen’s *d* = 2.01; Comp-Rot-Exp-Delayed group: 6.01 ± 2.47, t(11) = 8.43, *p* < 0.001, 95% CI = [4.44, 7.58], Cohen’s *d* = 2.43; fig.2A-olive and 2C-cyan).

Prior work has associated elevated reaction time with explicit mechanism engagement, reflecting the involvement of cognitive strategy during learning (Fernandez-Ruiz et al., 2011; Haith et al., 2015; McDougle et al., 2015; Taylor et al., 2014). Consistent with this, the transition from baseline to Session 1 (target-jump+gradual-rotation) showed a significant increase in reaction time across all groups. In immediate groups, reaction time increased significantly where we compared the reaction time at last bin of baseline with the first bin of learning session 1 (paired t-test, Comp-Rot-Imp-Imm group: t(11) = 2.42, *p* = 0.034, 95% CI = [8.56, 180.61], Cohen’s *d* = 0.87; Comp-Rot-Exp-Imm group: t(11) = 4.13, *p* = 0.0017, 95% CI = [74.07, 243.50], Cohen’s *d* = 0.99), indicating engagement of explicit strategy when learning the target jump (fig.2B-violet and 2D-pink). For Comp-Rot-Imp-Delayed group (fig.2B-olive), the increase in reaction time followed the same pattern and demonstrated a trend toward significance (paired t-test: t(11) = 2.11, *p* = 0.058, 95% CI = [−3.01, 147.18], Cohen’s *d* = 0.52). In Comp-Rot-Exp-Delayed group, the transition from baseline to Session 1 showed a significant increase in reaction time (fig.2D-cyan; paired t-test: t(11) = 3.01, *p* = 0.012, 95% CI = [34.80, 224.51], Cohen’s *d* = 0.79). This indicates that learning the target jump component reliably elicited explicit processing demands across participants during session 1.

These findings suggest that the simultaneous engagement of implicit and explicit mechanisms produces a compound adaptive state during learning. Both systems are active, and their outputs are integrated into a unified motor behavior.

### The compound memory is inflexible immediately after learning

As learning progressed during Session 1, reaction times gradually declined toward baseline, indicating that the explicit strategy became more efficient over trials. Critically, the transition from the compound learning block (Session 1) to the rotation-only block (Session 2) – in which the target jump was removed while the rotation persisted – revealed a striking carryover of the compound control policy. Despite the removal of the explicit demand, reaction times did not decrease; there was no significant difference between the last bin of Session 1 and the first bin of Session 2 in any of the four groups (fig. 2B, D). A two-way mixed ANOVA with factors Group (four levels) and Session (last bin of Session 1 vs. first bin of Session 2) confirmed no main effect of Session F(1, 44) = 0.00, *p* = 0.971, η²p < 0.001, no main effect of Group, F(3, 44) = 0.36, *p* = 0.783, η²p = 0.02, and no Group × Session interaction, F(3, 44) = 0.04, *p* = 0.989, η²p = 0.003.

Hand angle data provided convergent evidence for inflexibility. If the system had flexibly decomposed the compound memory, removal of the target jump should have produced an immediate shift to pure implicit adaptation – that is, hand angles near +20° (the full compensation required for the rotation alone). Instead, participants’ hand angles during the rotation-only session remained near the compound learning endpoint (approximately +10°). A paired-sample t-test comparing the average hand angle in the first bin of the rotation-only session (mean = 11.04 ± 0.32) against the pure implicit adaptation level during the last bin of Session 1 catch trials (mean = 5.51 ± 0.41) revealed a significant difference (t(47) = 12.82, *p* < 0.001, 95% CI = [4.66, 6.40], Cohen’s *d* = 2.20), confirming that behavior was not purely implicit. Moreover, when the contribution of the explicit target jump (∼27°) was removed from the compound learning hand angles, the resulting values did not differ from the subsequent rotation-only performance. A mixed ANOVA revealed no main effect of Sessions, F(1, 44) = 1.22, p = 0.275, η²ₚ = 0.03, indicating that behavior in Session 2 was statistically indistinguishable from the compound state. This suggests that the underlying implicit policy persisted unchanged across sessions.

Together, these results demonstrate that immediately after learning, the compound memory is inflexible: it cannot be rapidly decomposed into its constituent implicit and explicit components. Instead, the motor system continues to execute an integrated control policy, even when task demands no longer require both mechanisms. This pre-consolidation rigidity provides a critical baseline against which the effects of consolidation can be assessed.

### Immediate retention preserves the integrated compound memory

To assess whether the compound memory is expressed as an integrated whole immediately after learning, we compared hand angles from the last bin of the relevant learning session with those from the corresponding clamp trials. For the group that would later be probed on the implicit component, Comp-Rot-Imp-Imm, we compared hand angles at the end of the rotation-only session (the last block before retention) with those in zero-clamp retention trials. For the explicit probe group, Comp-Rot-Exp-Imm, we compared hand angles at the end of the compound learning block (jump + rotation) with those in jump-clamp retention trials. In both cases, hand angles during error-clamp trials did not differ significantly from the respective end-of-learning performances (Implicit probe: t(11)=1.8, *p* =0.099, 95% CI = [−0.28, 2.81], Cohen’s *d*=0.68; Explicit probe: t(11)=0.75, *p* =0.465, 95% CI = [−1.6, 3.28], Cohen’s *d*=0.24; fig. 2A, C, violet and pink bars). These results confirm that immediately after learning, the motor system retains the full compound policy: when cued to reproduce the rotation-only behavior (zero-clamp) or the target-jump+gradual-rotation behavior (jump-clamp), participants reproduced the integrated state they had just experienced.

### Consolidation enables task-dependent, component-selective expression

In order to assess whether consolidation separates the implicit and explicit components, and expression after 24 hours would reflect the task-dependent memory component, we compared the hand angles in the last bin of the learning session on Day-1 with those in the corresponding error-clamp trials after consolidation on Day-2. In both the implicit-probe delayed group (Comp-Rot-Imp-Delayed) and the explicit-probe delayed group (Comp-Rot-Exp-Delayed), the retention expression deviated significantly from the corresponding end-of-learning state (Comp-Rot-Imp-Delayed: t(11)=8.85, *p* <0.001, 95% CI = 1.27, 6.15], Cohen’s *d*=3.71; Comp-Rot-Exp-Delayed: t(11)=7.26, *p* <0.001, 95% CI = [5.82, 10.88], Cohen’s *d*=2.27). Direct comparison between immediate and delayed groups further underscored this reorganization: for the implicit probe, immediate vs. delayed zero-clamp hand angles were significantly different (t(22)=8.08, *p* <0.001, 95% CI = [6.21, 10.51], Cohen’s *d*=3.30); for the explicit probe, immediate vs. delayed jump-clamp hand angles also differed (t(22)=3.91, *p* <0.001, 95% CI = [3.07, 9.98], Cohen’s *d*=1.60). This suggests that consolidation does not merely preserve the compound memory; it actively alters the nature of expression.

To further investigate post-consolidation expression, we compared the delayed groups’ clamp-trial hand angles with the implicit aftereffects measured during compound learning (catch trials). For Comp-Rot-Imp-Delayed (zero-clamp, cued to reproduce rotation-only behavior), the expressed hand angles closely matched the implicit aftereffect from the compound learning catch trials (mean difference =0.9, t(11)=0.85, *p* =0.412, 95% CI = [−1.42, 3.22], Cohen’s *d*=0.28; fig. 2A, olive bars). This indicates that when the task context cues the rotation component, consolidation enables selective expression of the implicit component alone, without the explicit strategy. For Comp-Rot-Exp-Delayed (jump-clamp, cued to reproduce the target-jump+gradual-rotation behavior), the expressed hand angles did not match the compound behavior from the end of learning (t(11)=7.26, *p* <0.001, 95% CI = [5.82, 10.88], Cohen’s *d*=2.28). Instead, they closely resembled what would be expected if participants expressed primarily the explicit component, i.e., a hand angle near the explicit target-jump aim (approximately –30°) (fig. 2C, cyan bars). Thus, when the task context cues the jump component, consolidation enables selective expression of the explicit component, suppressing the implicit contribution. Overall, these findings indicate that post-consolidation motor behavior is task-dependent, selectively expressing the component of learning-implicit or explicit, that is most relevant to current task demands, rather than simply reproducing the original compound memory.

### Relearning reflects the component expressed at retention

The expression in the immediate groups showed the integrated compound memory, whereas the delayed groups expressed task-dependent, component-specific memories (implicit for the rotation cue, explicit for the jump cue). If consolidation truly separates components, then the memory expressed at retention should determine how quickly participants re-learn when exposed to the same perturbation. Re-learning should be faster when the expressed component matches the perturbation and slower when it mismatches (interference).

To test this, we examined relearning performance separately for the two probe types. For groups probed on the implicit component (rotation cue), relearning involved an abrupt –20° rotation with no target jump. The immediate group (Comp-Rot-Imp-Imm), which had expressed the full compound memory at retention, showed faster relearning than the delayed group (Comp-Rot-Imp-Delayed), which had expressed only the implicit component (t(22)=2.41, *p* =0.024, 95% CI = [0.26, 3.51], Cohen’s *d*=0.98; fig. 2A, relearning bins). This advantage indicates that the compound memory formed during initial learning remained active immediately after expression, enabling rapid reacquisition of the previously learned rotation. After 24 hours, however, the delayed group exhibited slower relearning, consistent with the shift toward a predominantly implicit state observed during their expression trials (fig.2A).

For groups probed on the explicit component (jump cue), relearning involved the –30° target jump alone (no rotation). Here, the pattern reversed: the immediate group (Comp-Rot-Exp-Imm), which had expressed the compound memory, relearned the jump more slowly than the delayed group (Comp-Rot-Exp-Delayed), which had expressed primarily the explicit component (t(22)=2.44, *p* =0.022, 95% CI = [0.65, 7.93], Cohen’s *d*=0.99; fig. 2C, relearning bins). The delayed group had an explicit strategy ready to use, so they learned the jump task quickly. The immediate group, however, still carried the full compound memory. That memory included an implicit component, which was not needed for the jump task and interfered with explicit aiming.

This pattern is further supported by the aftereffects observed in the jump-only condition. In the immediate group (Comp-Rot-Exp-Imm group), hand angles during catch trials were significantly different from zero (t(11) = 4.11, *p* = 0.0017, 95% CI = [1.67, 5.52], Cohen’s *d* = 1.19), indicating the implicit part of the compound mechanism which was still active (fig.2c). In contrast, the delayed group (Comp-Rot-Exp-Delayed group) showed no measurable aftereffects (t(11) = −0.30, *p* = 0.767, 95% CI = [−1.92, 1.43], Cohen’s *d* = −0.09), suggesting that consolidation had shifted behavior toward a predominantly explicit strategy with minimal implicit contribution (fig.2C-cyan).

Overall, our results from Experiment 1 show that simultaneous engagement of explicit and implicit processes forms a compound memory whose immediate expression reflects a unified inflexible memory, but consolidation reorganizes this memory into task-dependent expressions of its individual components, resulting in distinct patterns of expression and relearning in the immediate versus delayed conditions.

### Implicit mechanism is expressed immediately and persists after consolidation

Experiment 1 demonstrated that consolidation transforms a compound implicit-explicit memory into separable components. However, all participants in Experiment 1 who were later tested on the implicit component first completed a 100-trial rotation-only session before the retention test. This block was necessary to show that the initial memory was inflexible, but it also provided additional implicit practice. Therefore, the implicit aftereffect we observed after 24 hours could reflect the decay of the new implicit memory formed during that additional practice, rather than the original implicit component from compound learning. In Experiment 2, participants learned the exact same compound task (target-jump+gradual-rotation), but we removed the intervening rotation-only session. Immediately after learning, or after a 24-hour delay, we tested their implicit memory by asking them to simply “Target will not jump, aim straight to the target”. This allowed us to isolate the stability of the implicit aftereffect that remained from the original compound learning.

Participants from both groups successfully adapted to both perturbations simultaneously (last bin, Comp-NoRot-Imp-Imm group: –11.63 ± 2.31; Comp-NoRot-Imp-Delayed group: –12.19 ± 1.54). Similar to experiment-1, we observed significant aftereffects during catch trials of target-jump+gradual-rotation session (one sample t-test, Comp-NoRot-Imp-Imm group: 4.89 ± 2.57, t(11) = 6.59, *p* < 0.001, 95% CI = [3.26, 6.53], Cohen’s *d* = 1.9; Comp-NoRot-Imp-Delayed group: 3.59 ± 1.92, t(11) = 6.46, *p* < 0.001, 95% CI = [2.37, 4.82], Cohen’s *d* = 1.86), indicating the engagement of implicit mechanism (fig.3A). Further, the reaction time analysis revealed a significant main effect of Session, F(1, 22) = 7.93, *p* = 0.010, η²p = 0.081, indicating increased RT from baseline to the target-jump+gradual-rotation condition. There was no main effect of Group, F(1, 22) = 0.23, *p* = 0.638, η²p = 0.008, and no Group × Condition interaction, F(1, 22) = 0.31, *p* = 0.584, η²p = 0.003 (fig.3B). These results suggest that both implicit and explicit mechanism was active while learning target-jump+gradual-rotation session resulting in a compound mechanism engagement.

**Figure 3.**
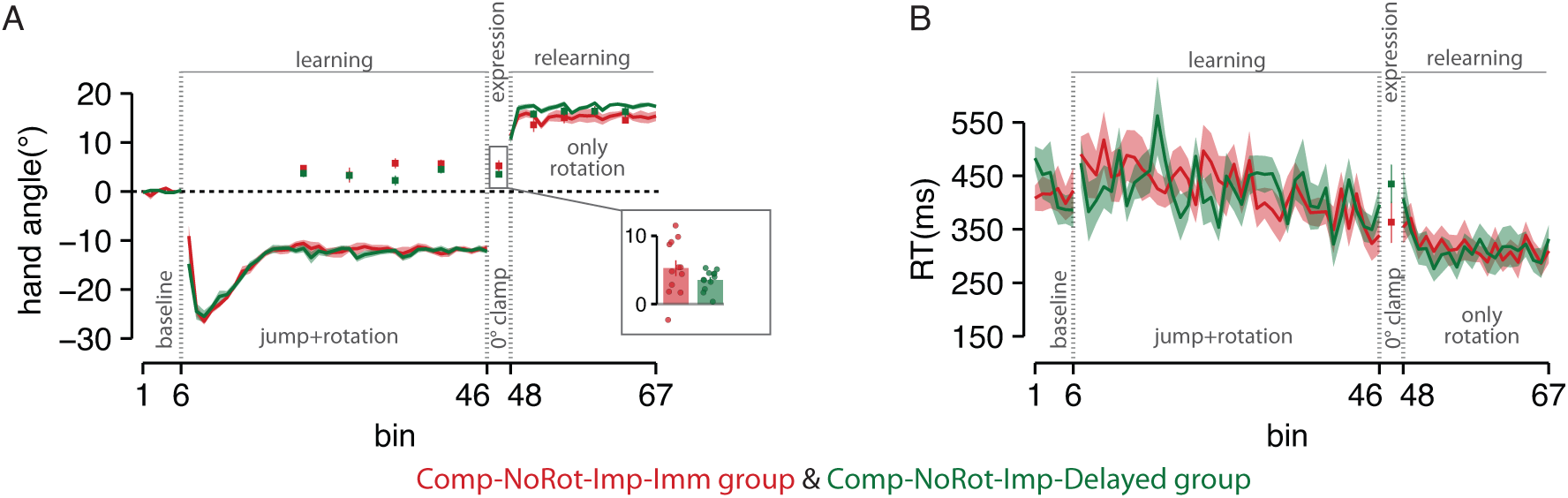
Implicit adaptation persists immediately and after 24-hour consolidation. A. Group-averaged, binned hand-angle trajectories during baseline, compound learning (target-jump+gradual-rotation), zero-clamp retention, and abrupt-rotation relearning. A. shows the Comp-NoRot-Imp-Imm group (red), and the Comp-NoRot-Imp-Delayed group (green). Both groups exhibited smooth adaptation to the compound perturbation and robust aftereffects during catch and zero-clamp trials, indicating engagement and persistence of implicit recalibration. The inset bar plot in A comparing the aftereffect during the retention block confirms that implicit aftereffects are comparable between immediate and delayed conditions. The relearning performance was comparable across both groups. B. Group-averaged, binned reaction-time dynamics for the immediate (red) and delayed (green) groups. RT increased during compound learning relative to baseline, reflecting explicit strategy engagement, but returned toward baseline levels during the zero-clamp retention phase when participants were instructed to aim directly at the target. Across groups, implicit aftereffects expressed during zero-clamp trials were comparable to catch-trial aftereffects from compound learning, demonstrating that implicit adaptation is expressed immediately and remains stable after 24-hour consolidation.

We examined aftereffects observed in the zero-clamped trials following compound learning to test the persistence of implicit learning, assessed immediately or 24 hours later. A one-sample t-test comparing hand-angle deviations during zero-clamp trials revealed significant aftereffects in both groups (Comp-NoRot-Imp-Imm group: 5.21 ± 3.96°, t(11) = 4.55, *p* < 0.001, 95% CI = [2.69, 7.73], Cohen’s *d* = 1.31; Comp-NoRot-Imp-Delayed group: 3.49 ± 1.45°, t(11) = 8.30, *p* < 0.001, 95% CI = [2.56, 4.42], Cohen’s *d* = 2.39). These results indicate that the implicit mechanism remained active during the expression phase. To further assess whether the aftereffects during zero-clamp trials reflect the same implicit component acquired during the compound learning session, we compared hand angles from the zero-clamp retention trials with those from the catch trials taken at the end of compound learning. A paired t-test showed no significant difference in Comp-NoRot-Imp-Imm group (t(11) = –0.51, *p* = 0.619, 95% CI = [-2.47, 1.54], Cohen’s *d* = – 0.13); the same pattern was observed after a 24-hour delay in Comp-NoRot-Imp-Delayed group (t(11) = –1.76, *p* = 0.105, 95% CI = [-2.22, 0.24], Cohen’s *d* = –0.35). This indicates that the implicit adaptation expressed during zero-clamp is consistent with the implicit component acquired during compound learning, regardless of whether it is expressed immediately or after consolidation (fig.3A). Additionally, we compared the magnitude of zero-clamp aftereffects across the two groups and found no significant difference between immediate and delayed expression (two-sample t-test: t(22) = 1.41, *p* = 0.172, 95% CI = [-0.81, 4.25], Cohen’s *d* = 0.57). Together, these results suggest that implicit adaptation acquired during compound learning is stable and doesn’t decay over the consolidation period.

After retention, when abrupt rotation was provided during the relearning session in Experiment 2, the immediate group (Comp-NoRot-Imp-Imm group) and the delayed group (Comp-NoRot-Imp-Delayed group) showed comparable relearning performance (fig.3A), with no reliable difference between the two conditions (two-sample t-test: t(22) = 0.65, *p* = 0.519, 95% CI = [−1.55, 2.98], Cohen’s *d* = 0.26). As both groups expressed the same state during expression, they started relearning from nearly identical motor states, yielding comparable relearning performance.

Overall, Experiment 2 demonstrated that the implicit component of the compound memory is both robust and stable: it can be expressed immediately after learning, persists across 24-hour consolidation, and remains unaffected by the absence of explicit strategy demands.

We have shown that simultaneous implicit-explicit learning produces a compound memory which later consolidates into separable components (Experiments 1 and 2). We next asked whether the order of learning changes this reorganization. In Experiment 3, participants first stabilized implicit adaptation to a gradual rotation in isolation before the explicit target-jump strategy was introduced. This design allowed us to test three specific predictions. First, if prior stabilization strengthens implicit learning, the implicit aftereffect measured during subsequent compound training should be larger than what we observed in Experiment 1 (where both mechanisms were learned simultaneously). Second, if the implicit system remains updatable, superimposing an explicit strategy onto an already-adapted implicit system should generate new sensory prediction errors, causing the implicit component to update. Third, and most critically, we asked which version of implicit memory consolidates: the original implicit-only state (cued by the rotation-only session) or the updated implicit state (measured during compound learning).

### The compound memory had stronger implicit aftereffects when initiated with implicit adaptation

Participants from all 4 groups implicitly adapted to the gradual perturbation, showing a slow, continuous recalibration of their hand movements without relying on strategic re-aiming. By the end of Session 1, participants exhibited a stable compensatory shift in hand angle consistent with implicit adaptation to the 20° rotation (last bin of session 1, mean ± SD = Rot-Comp-Imp-Imm group: 18.01 ± 1.37; Rot-Comp-Exp-Imm group: 17.62 ± 1.47; Rot-Comp-Imp-Delayed group: 17.58 ± 2.24; Rot-Comp-Exp-Delayed group: 16.81 ± 3.08). Following the rotation-only session, the introduction of the target jump on top of the previously acquired gradual rotation produced a reliable increase in reaction time (RT), indicative of explicit strategy engagement. A mixed ANOVA revealed a significant main effect of session (F(1,43) = 13.47, *p* < 0.001, η²p = 0.24), confirming that RTs increased when participants transitioned from rotation only session to the target-jump+gradual-rotation session. This suggests that the target jump successfully recruited explicit re-aiming even when implicit learning had already stabilized.

To quantify implicit adaptation during the target-jump+gradual-rotation session, catch trials were introduced in which participants were explicitly instructed that the target would not jump and asked to aim directly at the original target location. Across all four groups, hand-angle deviations during these catch trials were significantly different from zero, indicating the continued engagement of the implicit mechanism even while participants were concurrently using explicit strategies to counter the target jump (fig.4A and 4C). One-sample t-tests confirmed significant aftereffects in all groups (Rot-Comp-Imp-Imm group: 13.18 ± 2.06, t(11) = 22.09, *p* < 0.001, 95% CI = [11.87, 14.5], Cohen’s *d* = 6.37; Rot-Comp-Exp-Imm group: 11.97 ± 4.16, t(10) = 12.65, *p* < 0.001, 95% CI = [8.81, 12.57], Cohen’s *d* = 3.81; Rot-Comp-Imp-Delayed group: 12.02 ± 2.4, t(11) = 17.34, *p* < 0.001, 95% CI = [10.5, 13.55], Cohen’s *d* = 5.01; Rot-Comp-Exp-Delayed group: 11.97 ± 4.16, t(11) = 9.95, *p* < 0.001, 95% CI = [9.32, 14.62], Cohen’s *d* = 2.87), demonstrating that implicit adaptation remained robust and measurable in Session 2 despite the additional demands imposed by the explicit, target-jump–driven strategy. Critically, the magnitude of these aftereffects (mean = 11.97) was substantially larger than the implicit aftereffects observed in Experiment 1 (mean = 4.99), where implicit and explicit were learned simultaneously. A one-way ANOVA comparing the aftereffects from Experiment 1 and Experiment 3 confirmed a robust effect of experiment (F(1,93) = 152.50, *p* < 0.001, η²p = 0.62), with Tukey-adjusted post-hoc tests showing a significant increase in Experiment 3 (p < 0.001). This demonstrates that learning history could alter the component strength.

**Figure 4.**
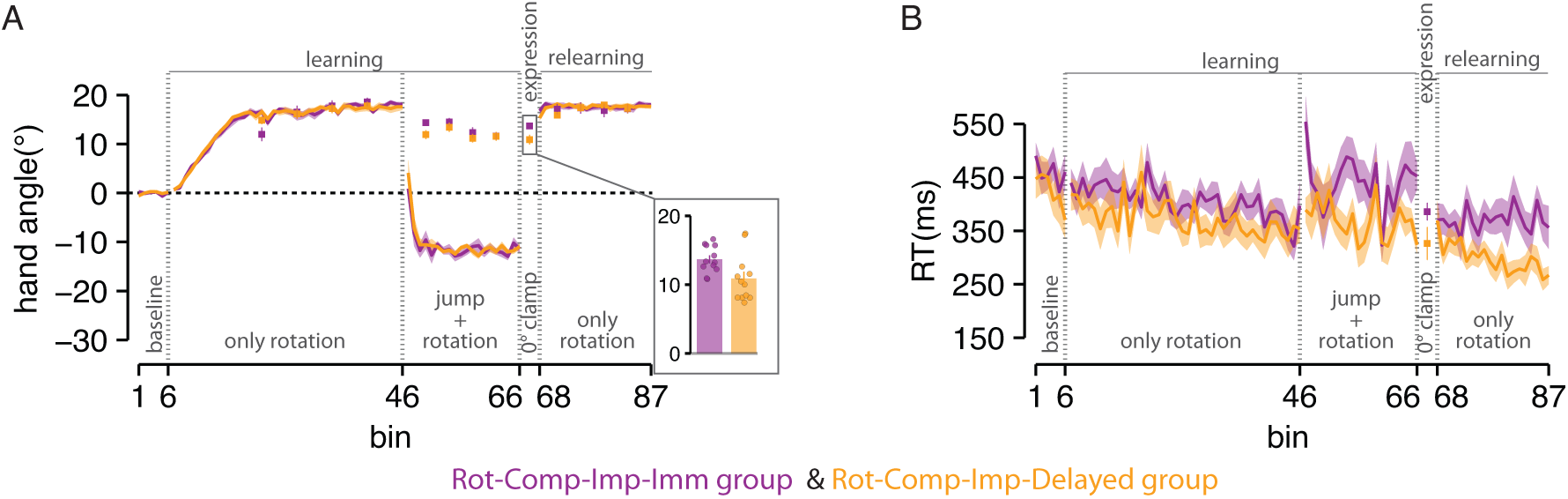
Expression and relearning after implicit-first learning exhibit task-driven implicit dominance. A, C. Group-averaged, binned hand-angle trajectories relative to the original target. Participants first showed robust implicit adaptation during the gradual-rotation block, followed by compound learning (target-jump + rotation), where implicit recalibration remained active alongside explicit strategy use, resulting in larger implicit aftereffects than in Experiment 1. Immediate groups (Rot-Comp-Imp-Imm, Rot-Comp-Exp-Imm) expressed an intermediate state between the original implicit-only adaptation and the updated implicit component (A, purple) and compound memory when the expression task required a jump response (C, blue). In contrast, after a 24-hour delay, delayed groups expressed the recalibrated implicit component when cued for rotation (A, yellow) and the explicit target-jump component when cued for jump (C, orange), indicating consolidation of the updated implicit state and component-selective retrieval. Inset bar plots (lower panels in A, C) compare expression performance between the immediate and delayed groups. Bar plots show that consolidation preserves the updated implicit component and promotes selective expression of implicit or explicit processes. The top inset panel in the C shows the after effects observed in the jump-only relearning session. This demonstrates that implicit aftereffects are present in the immediate group but disappear after consolidation, indicating a shift toward explicit-dominant expression in the delayed group during the jump-only relearning condition. B, D. Reaction times (group-averaged, binned) increased during introduction of the target jump, confirming explicit strategy engagement, and remained elevated in conditions requiring explicit control across both immediate and delayed groups.

Notably, the implicit aftereffect measured during the compound session of Experiment 3 (mean = 11.99) was smaller (t(46) = 9.02, *p* < 0.001, 95% CI = [3.15, 4.97], Cohen’s *d* = 1.47) than the original rotation-only adaptation (mean = 16.05). This reduction does not indicate decay or failure of updating. Instead, it reflects context-dependent recalibration: the addition of the target jump altered the sensory prediction errors that drive implicit learning, causing the implicit system to settle at a new, lower equilibrium. Critically, consolidation preserved this updated state (as shown by the delayed group), confirming that the change was a genuine memory update rather than transient suppression.

### Immediate expression reflects an updated state modulated by task demands

To assess immediate expression, we compared hand angles in the last bin of the learning session with those in the corresponding error-clamp trials. In the Rot-Comp-Imp-Imm group, where the task required participants to express only rotation behavior, participants’ hand angle was significantly different from the implicit-only behavior (fig.4A-purple). Instead, their expression fell between the original implicit-only state and the updated implicit aftereffect measured during Session 2 catch trials. Specifically, the zero-clamp hand angle was significantly different from the Session-2 catch aftereffect (t(11) = 3.26, *p* = 0.007, 95% CI = [0.67, 3.46], Cohen’s *d* = 0.91), indicating that immediately after learning, the memory was already shifting toward the updated state but had not fully resolved (fig.4A-purple). In contrast, for the Rot-Comp-Exp-Imm group (fig.4C-blue), when the task required immediate expression of a target-jump strategy, participants displayed behavior consistent with the engagement of the previously learned compound memory (paired t-test: t(10) = 1.23, *p* = 0.246, 95% CI = [-1.40, 4.86], Cohen’s *d* = 0.31).

### Consolidation preserves the updated implicit state and enables component-selective expression

To investigate whether consolidation preserved the updated implicit memory during the compound learning or the original one from the rotation-only session, we examined the expression in the Rot-Comp-Imp-Delayed group, where participants were asked to reproduce the rotation-only behavior. Their hand angles were significantly different from the original rotation-only state on Day-1 (paired t-test: t(11) = 5.17, *p* < 0.001, 95% CI = [3.84, 9.54], Cohen’s *d* = 2.32). Critically, their expression was not significantly different from the updated implicit aftereffect measured during Session 2 catch trials on Day-1(paired t-test: t(11) = –0.84, *p* = 0.418, 95% CI = [-4.11, 1.83], Cohen’s *d* = –0.38). This indicates that consolidation stabilized the updated implicit component

Next, we asked whether consolidation reorganized the compound memory into its components, and explicit memory was expressed when the task demanded, despite the initial stabilization of the implicit memory on Day 1. Participants in the Rot-Comp-Exp-Delayed group during the jump-clamped trials did not reproduce the compound behavior from initial learning; rather, their performance was biased toward the explicit target-jump strategy (–30 degree), similar to the expression observed previously in the experiment 1 (Comp-Rot-Exp-Delayed group) (fig.4C-orange). Further, the difference between the explicit immediate and delayed groups approached significance (t(21) = 2.04, *p* = 0.053, 95% CI = [-0.07, 10.02], Cohen’s *d* = 0.85). This indicates that, under target-jump task demands, consolidated memory selectively expresses the individual component over the compound memory.

### Relearning reflects the expressed component during retention

Despite the differences in retention expression observed earlier – immediate groups expressed a hybrid state, whereas delayed groups expressed the pure updated implicit state – the two groups relearned at comparable rates. A two-sample t-test revealed no significant difference between the Rot-Comp-Imp-Imm and Rot-Comp-Imp-Delayed groups (t(22) = 1.64, *p* = 0.114, 95% CI = [-0.37, 3.26], Cohen’s *d* = 0.67; fig. 4A). This suggests that consolidation did not differentially reorganize or bias the expressed memory in this condition. In contrast, in the target-jump only relearning session, the Rot-Comp-Exp-Imm group, which had expressed the full compound memory at retention, relearned the jump task significantly more slowly than the Rot-Comp-Exp-Delayed group, which had expressed a primarily explicit strategy (t(21) = 3.50, *p* = 0.002, 95% CI = [2.04, 8.01], Cohen’s *d* = 1.46; fig. 4C). This pattern mirrors Experiment 1: carrying forward a compound memory that includes an implicit component interferes with relearning a pure explicit task, whereas having already separated the explicit component facilitates rapid reacquisition. The relearning differences were further supported by the aftereffects measured under the jump-only relearning condition. Rot-Comp-Exp-Imm group exhibited significant aftereffects (one sample t-test: t(10) = 3.87, *p* = 0.003, 95% CI = [1.71, 6.37], Cohen’s *d* = 1.16), indicating continued expression of the implicit component of the compound memory (fig.4C-blue). However, Rot-Comp-Exp-Delayed group showed no reliable aftereffects (one sample t-test: t(11) = 1.20, *p* = 0.254, 95% CI = [−1.48, 5.04], Cohen’s *d* = 0.34), reflecting a shift toward an explicitly driven state with negligible implicit contribution (fig.4C-orange). Together, these results demonstrate that immediate groups retain access to a compound memory, whereas delayed groups express selectively implicit or explicit components, leading to distinct relearning dynamics across tasks.

## Discussion

Our study provides insights into how consolidation transforms compound motor memories formed through simultaneous engagement of implicit and explicit learning mechanisms. Across three experiments, we show that engaging implicit (gradual rotation-driven) and explicit (target jump-driven) mechanisms simultaneously creates a compound adaptive memory whose expression is initially unified. However, consolidation transforms this compound memory into its implicit and explicit components, making them selectively expressible depending on task demands. Moreover, when implicit learning is allowed to stabilize before explicit demands are introduced, the implicit component becomes substantially stronger and remains updatable, and consolidation preserves this updated state even though it is quantitatively weaker than the original implicit-only memory. These findings challenge a purely stabilization view of consolidation and instead suggest that consolidation actively restructures motor memories into flexible, context-sensitive components.

### From immediate integration to delayed selectivity: how consolidation restructures memory

Immediately after learning, the motor system treated the compound memory as a single, unified whole. Participants expressed the compound motor output despite the task demands of rotation or target-jump behavior. This pattern indicates that the system did not store two independent memories (one implicit, one explicit) during the learning. Rather, it formed a single, integrated memory state that contained the computations for both behaviors. The immediate retention test, therefore, reveals the pre-consolidation buffer state, which has an active internal model that blends both learning mechanisms, not a passive repository of separate representations (Krakauer et al., 2019; Smith et al., 2006). This observation aligns with recent work showing that implicit and explicit processes interact competitively during learning (Albert et al., 2022; McDougle et al., 2015), but it extends those findings by demonstrating that the interaction produces a functional integration that persists even when one of the task demands is removed (A. D. Kumar et al., 2025). Indeed, using a paradigm that drove learning in specific task dimensions, Miyamoto, Wang, and Smith (2020) found that strategy and implicit adaptation synergize in driven dimensions. Critically, they showed that the implicit system learns to compensate for the noise and variability introduced by an erratic explicit strategy, effectively “cleaning up” the performance. This functional cooperation explains why, in our study, the explicit component could not be immediately separated: the two systems had already adapted to each other’s presence, creating a unified control policy.

The expression following a 24-hour delay represents a fundamental reorganization of memory structure rather than mere stabilization of existing representations. Instead of reproducing the compound behavior, consolidation led to the selective expression of components relevant to the task context. When cued to perform the rotation-alone task (zero-clamp), they expressed a pure implicit aftereffect of approximately 5°, matching the catch-trial aftereffect from compound learning. When cued to perform the jump-alone task (jump-clamp), they expressed the explicit strategy, and no measurable implicit aftereffect. This component-selective expression was completely absent immediately after learning. Thus, consolidation did not merely stabilise the existing memory; it actively reorganised the memory structure, transforming an integrated representation into two separable components whose expression could be gated by task context.

One possible explanation could be based on the contextual inference (COIN) framework, which proposes that the brain adaptively controls memory expression in response to environmental cues (Heald et al., 2021, 2023). Consolidation might reduce uncertainty about which latent context (implicit-only, explicit-only, or compound) generated the sensory outcomes during learning. Initially, both mechanisms are associated with the same task context, forming an integrated representation. Over time, consolidation strengthens the association between each component and its distinctive retrieval cue (rotation-alone instruction vs. jump-alone instruction), allowing task demands to selectively gate expression. This view predicts that after consolidation, the system can flexibly retrieve the appropriate component without interference.

### Learning history alters component strength and updatability

In Experiment 3, when implicit adaptation to a gradual rotation was stabilized in isolation, and then the target jump was added, aftereffects were more than twice as large as in Experiment 1 (13° vs. 5°). Prior stabilization dramatically strengthened the implicit component, indicating that the interaction between learning systems is not fixed but depends on the order and timing of experiences. This aligns with the finding that gradually introducing a visuomotor rotation leads to substantially larger implicit adaptation than abrupt introduction, as the brain integrates the error over time without triggering strategy-based compensatory mechanisms (Ruttle, 2023).

Moreover, this aftereffect was significantly smaller than the original rotation-only adaptation (13° vs. 18°). This reduction might be mistaken for decay or interference. However, it reflects context-dependent recalibration: adding the target jump changed the sensory prediction errors that drive implicit learning. This update is consistent with the idea that implicit and explicit systems compete for a common error (Albert et al., 2022). When the explicit system is engaged, it takes away part of the available error signal, thereby reducing the effective drive for implicit adaptation. Crucially, consolidation preserved this updated state. If the reduction were due to transient suppression, the original, stronger memory should have re-emerged after 24 hours. It did not. Thus, the implicit system remains malleable and continues to integrate ongoing experience even when explicit strategies are deployed (Bond & Taylor, 2015; Mazzoni & Krakauer, 2006; Shadmehr et al., 2010).

The finding that consolidation preserves the updated, weaker state aligns with evidence that implicit adaptation remains malleable after initial stabilization. Miyamoto, Wang, and Smith (2020) showed that the implicit system dynamically compensates for noise introduced by an explicit strategy, continuously recalibrating rather than freezing. Benson, Anguera, and Seidler (2011) found that an explicit spatial strategy modulates the progression of implicit recalibration, leading to a lower aftereffect. Avraham and colleagues (2021) demonstrated that implicit adaptation attenuates during relearning rather than simply re-expressing. In Experiment 3, the explicit re-aiming strategy temporarily reduced the error drive to the implicit system, causing it to settle at a new, lower equilibrium (13°). Consolidation then locked in this updated state rather than reverting to the original stronger state (18°). Thus, the implicit system is not rigid; it integrates ongoing experience, and consolidation preserves the most recent functional state regardless of whether it is stronger or weaker than a previous one.

### Neural basis of component separation and links to other memory systems

The reorganization from inflexible integration immediately after learning to task-selective expression after 24 hours strongly implicates systems consolidation, in which memories undergo time-dependent restructuring across distributed neural circuits (Dudai et al., 2015). Critically, implicit and explicit learning engage distinct but interacting neural systems. Implicit sensorimotor recalibration depends on cerebellar error-correction circuits that modify forward models (Smith & Shadmehr, 2005; Tseng et al., 2007), whereas explicit strategy use engages prefrontal and parietal networks involved in cognitive control (Doyon et al., 2009; Taylor et al., 2014). Our behavioural data suggest that during simultaneous learning, these systems are co-activated and their outputs are bound together into a unified representation. Over time, consolidation restructures this co-activated network, allowing the two components to be retrieved independently.

Neuroimaging studies of motor sequence learning provide direct evidence for such consolidation-dependent network reorganization. Using dynamic causal modelling, Tzvi and colleagues (2015) found that early motor memory encoding involves widespread modulation of cortico-striatal and cortico-cerebellar connections, whereas after consolidation, the network becomes pruned, with a more focused connection from left cerebellum to right putamen emerging as a key pathway. Similarly, consolidation alters sequence-specific distributed representations: representations in dorsolateral striatum, prefrontal cortex, and secondary motor cortex become stronger for consolidated sequences, while hippocampal and dorsomedial striatal representations become less engaged (Albouy et al., 2013). In our context, the initial compound memory may be supported by co-activation of cerebellar (implicit) and prefrontal (explicit) circuits. Consolidation then strengthens the independent retrievability of each pathway, perhaps by reducing cross-talk or establishing task-specific gating mechanisms. This interpretation aligns with the model proposed by Doyon and colleagues, in which different phases of motor learning – encoding, consolidation, and retention – are mediated by distinct but interacting cortico-striatal and cortico-cerebellar networks (Doyon et al., 2009, 2018).

Remarkably, this principle of consolidation-dependent segregation of initially integrated representations is not unique to motor learning. In declarative memory, the standard systems consolidation theory posits that new episodic memories initially depend on the hippocampus, but over time they become represented in neocortical networks independently of the hippocampus (Squire & Alvarez, 1995). More recent work has shown that consolidation involves a shift from hippocampal-centred retrieval networks to neocortical-centred ones, with posterior hippocampal activity decreasing and neocortical activity increasing as memories mature (Takashima et al., 2009). This parallels our finding that immediate expression reflects a unified, integrated state (akin to hippocampal-dependent episodic retrieval), whereas delayed expression reflects component-selective, task-dependent retrieval (akin to neocortical-dependent semantic retrieval). In both domains, consolidation transforms memory from a context-general, integrated representation to a context-sensitive, modular one. Moreover, Keisler and Shadmehr (2010) showed that a fast, declarative-like component of motor memory can be disrupted by a declarative memory task during consolidation, whereas a slow, implicit component is immune – a finding that mirrors the dissociation we observed after 24 hours. Taken together, these convergent findings point to a common principle: consolidation actively reorganizes memories, sacrificing immediate unity for later flexibility.

### Relearning depends on the expressed component, not simply prior experience

The relearning data provided the strongest evidence that consolidation changes memory structure rather than just performance. In Experiment 1, immediate groups (which expressed the full compound memory) showed faster relearning of the rotation but slower relearning of the jump compared to delayed groups (which expressed pure implicit or pure explicit components). This pattern directly follows from the match or mismatch between the expressed component and the relearning task. In Experiment 2, because both groups expressed the same implicit state, relearning was identical. In Experiment 3, the implicit probe groups showed comparable relearning (both expressed the updated implicit state), while the explicit probe groups showed a clear advantage for the delayed group, which had already separated the explicit component.

These findings have important implications for the concept of savings in motor adaptation. The traditional view of savings – faster relearning upon re-exposure to a previously experienced perturbation – has often been attributed to the retention and rapid re-expression of an internal model. However, recent work has shown that savings is not a unitary phenomenon. Avraham and colleagues (2021) found that explicit and implicit processes exhibit opposite effects on relearning: explicit learning shows clear savings, whereas implicit adaptation is attenuated. This aligns closely with our observations: in Experiment 1, the immediate explicit group (which retained the explicit component) showed faster jump relearning, while the delayed implicit group (which expressed only the implicit component) showed slower rotation relearning – consistent with attenuation rather than savings of the implicit process. Our results also speak to the debate about whether savings reflect model-based (explicit) or model-free (implicit) mechanisms. Huang and colleagues (2011) argued that savings is driven by model-free processes such as use-dependent plasticity and operant reinforcement, which are closely linked to explicit strategy use. The present findings support this view: savings occurred only when the expressed component matched the task – that is, when the explicit component was available and relevant. When relearning required implicit adaptation alone, savings were absent or reversed.

## Conclusion

The most striking finding of this study is that consolidation does not simply protect the strongest or most recent memory; it actively re-evaluates which version of a memory component is most appropriate for the current sensorimotor context. Even when the updated implicit state is quantitatively weaker than the original adaptation, consolidation preserves it. This reveals a principle of memory optimization: the brain sacrifices raw strength for contextual relevance. More broadly, we have shown that motor memories formed through simultaneous implicit and explicit learning are initially unified and inflexible. Overnight consolidation reorganizes them into dissociable components whose expression depends on task context. The temporal order of learning determines component strength, and the implicit system remains updatable even after stabilization. These findings advance our understanding of motor memory consolidation by showing that it involves active restructuring to create flexible, context-sensitive representations – not merely the strengthening or decay of a single trace. This principle may generalize to other forms of multi-system learning and has practical implications for rehabilitation, skill training, and human-machine interaction, where implicit and explicit processes must be balanced.

## CONFLICT OF INTEREST

None

## ACKNOWLEDGEMENT

We would like to thank IIT Hyderabad for all the institutional-level support.

## DATA AVAILABILITY

The datasets used and/or analyzed during the current study are available from the corresponding author on reasonable request.

## FUNDING

This work was supported by DST CSRI (DST/CSRI/2021/164(C)(G)) grant to NK.

## Notes

### Competing Interest Statement

The authors have declared no competing interest.

